# Fantastic beasts and how to sequence them: genomic approaches for obscure model organisms

**DOI:** 10.1101/165928

**Authors:** Mikhail V. Matz

## Abstract

Application of genomic approaches to “obscure model organisms” (OMOs), meaning species with little or no genomic resources, enables increasingly sophisticated studies of genomic basis of evolution, acclimatization and adaptation in real ecological contexts. Here, I highlight sequencing solutions and data handling techniques most suited for genomic analysis of OMOs.

**Glossary:**

- **Allele Frequency Spectrum, AFS** (same as Site Frequency Spectrum, SFS): histogram of the number of segregating variants depending on their frequency in one or more populations.
- **Restriction site-Associated DNA (RAD) sequencing**: family of diverse genotyping methods that sequence short fragments of the genome adjacent to recognition site(s) for specific restriction endonuclease(s).
- **Linkage Disequilibrium (LD)**: in this review, correlation of genotypes at a pair of markers across individuals.
- **LD block**: typical distance between markers in the genome across which their genotypes remain correlated.
- **Genome scan:** profiling of genotypes along the genome looking for unusual patterns. Often used to look for signatures of natural selection or introgression.
- **“Denser-than-LD” genotyping**: genotyping of several polymorphic markers per LD block.
- **Highly contiguous reference**: genome or transcriptome reference sequence containing the least amount of fragmentation.
- **Phased data**: data showing which SNP alleles belong to the same homologous chromosome copy.
- **Cross-tissue gene expression analysis**: looking for individual-specific shifts in gene expression detectable across multiple tissues. Such shifts are predominantly genetic in nature.

---

The focus of this review is mainly on the type of sequencing data required and how to obtain it in the most cost-efficient way rather than on analytical approaches. That said, I could not help but mention some highly promising analytical methods that are not yet broadly adopted by OMO researchers, such as demographic inference based on allele frequency spectra and annotation-independent analyses of gene expression data.

I will start with the summary of general types of questions in OMO studies and corresponding data types required. We might be interested in the following four layers of genomic information, each requiring a specific type of experimental and reference data:

1. Genome-wide patterns of neutral variation. This data can elucidate population structure, population sizes, and migration rates, as well as changes of these parameters through time. This analysis benefits from high quality genotype calls but does not require dense genome coverage; it can even be performed in the absence of reference genome.
2. Regions in the genome particularly affected by non-drift processes (natural selection, introgression, etc). This type of analysis, typically referred to as “genome scanning”, takes genome-wide neutral variation as baseline and looks for regions in the genome exhibiting highly dissimilar patterns. It requires “denser-than-LD” genotyping and a highly contiguous reference (see Glossary) to make sure no signal is overlooked.
3. Genome-wide gene expression, an extremely information-rich resource reflecting both environmental and genetic variation. Streamlined transcript counting methods represent a cost-efficient alternative to the industry-standard RNA-seq for generating quantitative data. Analysis of gene expression does not require a genome reference, although a transcriptome reference must be generated at some point. The reference does not have to be highly contiguous.
4. Epigenetics, here limited to DNA methylation. A variety of methods have been recently developed that can generate data for DNA methylation analysis. For vertebrates, genome reference is needed, but for other animals or plants, in which DNA methylation is much less prevalent and predominantly occurs in exons, transcriptome or exome presents a good cost-efficient alternative. The reference does not have to be highly contiguous.

## Genome-wide neutral variation

### Allele Frequency Spectrum analysis

Neutral genetic markers are traditionally analyzed using summaries of allele frequency differences between populations, such as *F*_ST_. The large amount of markers accessible through next-generation sequencing opened up the possibility to dramatically enhance this approach by modeling the evolution of the whole allele frequency spectrum (AFS, see Glossary). AFS represents a rich source of information to fit alternative models with time-resolved population sizes and migration rates as parameters (Box 1) based on coalescent simulations (*fastsimcoal2*, [1]), diffusion approximation (*dadi*, [2]), or ordinary differential equations (*moments*, [3]). Model selection is then based on likelihood ratio tests or Akaike information criterion. The new *moments* method is particularly promising, as it is substantially faster than its predecessors and includes built-in bootstrap, demographic model plotting, and capacity to analyze up to five populations simultaneously. It is also very helpful that *moments* inherits the python code structure well familiar to *dadi* practitioners.

### Experimental data

The data required for AFS analysis is several thousand biallelic neutral single nucleotide polymorphisms (SNPs). Ideally, SNPs must not be closely physically linked in the genome to represent independent data points, although it is fully appropriate to analyze linked SNPs with AFS methods. The lack of requirement for contiguous SNP coverage makes various flavors of restriction site-associated DNA (RAD) sequencing (recently reviewed in [4,5], see Glossary) well suited for this analysis. In our experience, *dadi* [2] and *moments* [3] work robustly with 5-10 thousand SNPs (a typical RAD output) when analyzing populations individually or in pairs. Fitting models with three (*dadi*) or more (*moments, fastsimcoal2*) populations might be problematic with this relatively low number of SNPs but is usually not required for OMOs (Box 1). Recent population size changes are often of special interest in OMOs; since they predominantly affect rare alleles, their robust detection requires 20 or more high-quality genotypes per population [6]. This preference for more individuals rather than more SNPs per individual is an additional factor that makes cost-efficient RAD the approach of choice for AFS-based analysis. That said, relatively low number of independent (unlinked) SNPs generated by some RAD protocols might limit the power of the AFS analysis, and a good subject for a future study would be the effect of the number of unlinked SNPs on AFS model selection and uncertainties of parameter estimates. In this regard it is worth noting that RAD flavors differ considerably in the number of unlinked loci in the genome that they interrogate [4,5].

For demographic inference, the AFS data must be filtered to exclude potential sites under selection. Whichever test is used to identify such sites (for example, Bayescan, [7]), for their removal the false discovery rate should be set as high as 0.5 to ensure purging of the majority of non-neutral sites. Although under this setting half of the removed sites would be neutral, their removal will not affect the overall AFS as long as the removed fraction does not comprise more than 1-2% of the total number of sites.

### Genotyping quality

In diploids, the most common genotyping error is missing one of the alleles in a heterozygote (i.e., a false homozygote call); and the next most common error is missing the whole SNP locus entirely. Both these “missing data” errors are due to insufficient sequencing coverage, the problem that is pervasive in today’s OMO studies. Such errors strongly affect AFS in the region of rare alleles, which is unfortunate since rare alleles are the most informative about recent population history [6,8]. A telltale sign of poor heterozygote calling is under-representation of singletons, but frequencies of doubletons and higher-order frequency bins are also distorted, which has strong effect on AFS itself and inferred demographic parameters until mean sequencing coverage approaches ∼10x [9]. When coverage is 10x or higher a good way to filter data is to select SNPs genotyped in >90-95% of samples [10]; importantly for RAD approach, this would select SNPs that are unlikely to be affected by null alleles due to mutation in the restriction endonuclease recognition site [4]. For obvious reasons, for AFS analysis genotype calls should never be quality-filtered based on allele frequencies (for example, retaining only variants that are detected in a minimum of two individuals or requiring minor allele frequency to exceed some cutoff). A robust empirical way to evaluate the consistency of genotype calls is to compare results for independently processed biological samples of the same genotype [11]. Such genotyping replicates are quite feasible in RAD and are also useful to identify true SNPs for training variant quality score recalibration model of the GATK pipeline [12]. For low-coverage data (<10x), a general solution is provided by the *ANGSD* package [13], which generates AFS as well as other population genetic statistics based on genotype likelihoods without actually calling genotypes [14]. This method generates unbiased single-population AFS even with 2x coverage [9]. Still, there is a concern that high variation in coverage across samples and populations might affect *ANGSD* statistics; to avoid this potential issue it is recommended to discard the lowest-coverage outliers and down-sample reads from highest-coverage outliers (J. Ross-Ibarra, pers. comm.).

### PCR duplicates

Presence of PCR duplicates in many early RAD applications has been repeatedly highlighted as a source of genotyping errors [4,15] due to induced over-dispersion of read counts among alleles and loci. Interestingly, the proportion of PCR duplicates does not depend on the number of PCR cycles performed during library preparation. Instead, it depends on the ratio between the number of reads sequenced (*N*_r_) and the number of unique fragments present in the sample prior to PCR (*N*_o_): the fraction of duplicates is the same as expected when sampling *N*_r_ from *N*_o_ with replacement. Fortunately, PCR duplicates are easy to identify and remove using degenerate tags ligated to RAD fragments prior to amplification [16]. Most present-day RAD protocols now implement this simple deduplication procedure, including the current version of 2bRAD [11].

### Genome reference for AFS analysis

A great advantage of RAD-based AFS analysis for OMOs is that SNPs can be called based on RAD reads themselves, without the need for genome reference. Several *de novo* RAD genotyping pipelines have been developed, such as STACKS, pyRAD, and UNEAK (see references in [4]) that work for most RAD flavors, plus a similarly structured pipeline for 2bRAD (https://github.com/z0on/2bRAD_denovo) that takes into account the fact that in 2bRAD either strand of the locus can be sequenced. Still, using a reference genome to call RAD genotypes provides three important advantages. First, it identifies physically linked (and thus potentially non-independent) groups of SNPs, to be resampled as units during AFS bootstrap. The second advantage is particularly important for OMOs sampled in the field: mapping to reference genome automatically discards reads from contaminant DNA sources (viruses, bacteria, ingested food, symbionts etc). To be able to discard such contaminants in *de novo* RAD pipeline the experiment must include at least one sample generated from a clean source and consider only the RAD loci observed in that sample.

The third advantage of reference-based genotyping is the possibility to discriminate between ancestral and derived SNP alleles, to attain the best power of AFS-based inference. Counter-intuitively, the best reference for AFS analysis is not a genome of the species under investigation but a genome of a related outgroup species, separated from the focal one by a few million years of evolution, because the SNP state as in the outgroup can be assumed to represent the ancestral state (e.g., [17]). If the reads are mapped to the same-species genome, to identify ancestral states of the variants a single well-sequenced RAD sample of an outgroup taxon could be included. The analysis will then be limited to sites that can be successfully genotyped both in ingroup and outgroup; in effect, the result is going to be the same as when mapping the reads from whole project to an outgroup’s genome. Although some proportion of ancestral states will be misidentified due to incomplete lineage sorting, convergence or technical artifacts, this error is easy to account for by including a single additional parameter into the model, specifying the proportion of the AFS that needs to be flipped when predicting the data (e.g., [18]). The reference for AFS does not have to be highly contiguous; the contigs should be just long enough to cover a typical LD block for meaningful bootstrapping.

## Genome scanning

Since outlier regions by definition occupy only a small portion of the genome and typically do not form a single cluster, their confident detection requires “denser-than-LD” genotyping (see Glossary). It has been argued that in most situations, RAD-like approaches would sample the genome too sparsely to satisfy this requirement [19,20]. Although many successful genome scans based on RAD have been published [21], RAD cannot be recommended for genome scanning since it inevitably leaves considerable fraction of the genome unexplored. Even when LD is known to be extensive enough for RAD to produce “denser-than-LD” genotyping, a better solution might be to take full advantage of the extended LD and go instead for low coverage whole-genome sequencing (WGS) followed by imputation, to obtain full-genome phased data (Table 1).

**Table 1.** **Genotyping approaches for genome scanning.**

| Approach | Features | Pros | Cons |
| --- | --- | --- | --- |
| Exome-seq<br>[48,49] | Isolates and sequences only the protein-coding portion of genome. | Dense coverage of genes guarantees that coding variants and variants linked to cis-regulatory mutations are discovered. | Other (arguably less important) types of variation are not profiled (e.g., distant enhancers). |
| RNA-seq<br>[50,51] | Sequences RNA. | Same as exome sequencing. | Genotyping quality of a gene depends on expression level. Allele-specific expression affects accuracy of heterozygote calls. |
| Pool-seq<br>[52,53] | Sequences pooled DNA from multiple individuals from each population. | Dense whole-genome coverage with confident determination of allele frequencies in populations. | No possibility for individual-based analysis (such as STRUCTURE) or validation based on genotype-phenotype association across individuals. Must be confident in *a priori* population designations. |
| Low-coverage whole-genome sequencing (WGS)<br>[54] | Sequences individual genomes at ~1-4x coverage. | Dense whole genome coverage at individual level. | Per-site genotypes are unreliable because of missing data; must use uncertainty-aware analysis such as *ANGSD*. |
| Ultra-low coverage WGS with imputation<br>[22] | Sequences individual genomes at <2x coverage, imputes missing genotypes and corrects false homozygote calls | Dense whole genome coverage at individual level, phased data enables haplotype-based analysis | Rare alleles (minor allele frequency<0.05) are missed. Requires large sample sizes (depending on LD, hundreds or thousands of individuals). Accuracy of imputation must be experimentally validated for every new OMO. |

The types of sequencing approaches for genome scanning with their pros and cons are summarized in Table 1. Importantly, all of them require highly accurate reads mapped to a reference for confident SNP detection, making short Illumina reads the genotyping data type of choice. Some of the very promising approaches that have not yet been fully adopted for OMOs are exome-seq and ultra-low whole genome sequencing (WGS) with imputation. Exome-seq used to be a prerogative of model organisms because of the need for exome-capture platform development, but it has recently been shown that OMO exome can be captured just as efficiently using bead-bound normalized cDNA obtained from the OMO itself (EecSeq Puritz 2017). Such “home-made exome” sequencing could become an excellent alternative to RAD since it would interrogate essentially all the interpretable genetic variation for a comparably low cost. Ultra-low WGS with imputation used to require extensive reference haplotype panels available only for well-established model organisms. However, several methods have been recently developed (most notably STITCH, [22]) that can impute phased genotypes and correct genotyping errors in ultra-low coverage data without relying on reference panels. Still, their applicability for each new OMO must be experimentally confirmed because the success of imputation critically depends on multiple polymorphisms occurring within a typical LD block, and whether this is so is not known for OMOs *a priori*. Demographic events such as strong recent bottleneck, domestication, or recent colonization would make imputation more efficient because of more extensive LD and small number of founding haplotypes [22], and conversely, in large outbred populations imputation will be less accurate and might require sequencing of a very large number (thousands) of individuals. The accuracy of imputation can be evaluated by sequencing a few individuals at high coverage (>10x) to generate high-confidence genotype calls and then attempting to impute them based on sub-sampled read sets to emulate low coverage. It must be noted that it is inappropriate to measure imputation accuracy by imputing genotype calls masked in high-quality datasets (as in, for example, [23]): masked data do not contain false homozygote calls and therefore do not correctly represent the real-life situation.

## Gene expression

There are many aspects to gene expression, of which I here focus on just one: abundance or protein-coding (polyadenylated) transcripts. The reason is that transcript abundance is by far the most interpretable and it can be very easily analyzed in OMOs.

### Counting transcripts instead of resequencing them

Typical RNA-seq [24] resequences the whole transcriptome in each sample, but there is a much more economic way to count abundances of protein-coding transcripts: sequence just a single fragment per each transcript molecule and count reads corresponding to each gene. TagSeq [25], for example, sequences a single randomly generated fragment near the 3’-end of the transcript, which is the most economic use of sequencing effort and removes bias towards longer transcripts. In a recent benchmarking study TagSeq was actually more accurate than the standard RNA-seq in measuring transcript abundances, despite nearly tenfold lower cost [26]. More recently introduced QuantSeq [27] is conceptually similar to TagsSeq: it also sequences a single randomly generated fragment near the 3’-end of each transcript but has a different library preparation procedure, implemented as a kit from Lexogen (https://www.lexogen.com/quantseq-3mrna-sequencing/). Bioinformatics analysis for both TagSeq ad QuantSeq is highly simplified compared to typical RNA-seq. TagSeq was originally designed for OMOs and so its pipeline uses transcriptome rather than genome as a reference to attribute reads to genes (https://github.com/z0on/tag-based_RNAseq). One notable feature of the current version of TagSeq pipeline is that it includes removal of PCR duplicates based on adaptor-derived degenerate tags [11], similarly to 2bRAD and for the same reason – to avoid PCR-associated over-dispersion or read counts.

### Analysis of gene expression “beyond gene lists”

The unfortunate tradition that OMO research inherits from the biomedical field is putting too much emphasis on possible functional implications of expression changes of specific genes. For OMOs, this is bound to remain inconclusive because gene annotations are often absent, tentative or based predominantly on similarity to human genes, which may or may not serve the same function in the OMO. Even greater problem is interpretation bias: too often researchers focus primarily on genes that “make sense” and ignore the rest. This leads to conclusions reflecting predominantly the researchers’ idea of what *should* be going on rather than what is actually happening.

Table 2 lists alternative ways of objective analysis of gene expression data that are enabled by the large sample sizes feasible with TagSeq or QuantSeq. They either do not require gene annotations or rely sufficiently general functional summaries to be robust to occasional missing or mis-annotations. Particularly useful for OMOs are analyses that use gene expression patterns as anonymous multivariate readouts to compare and classify samples, such as principal coordinate analysis (PCoA) or differential analysis of principal components (DAPC). Related multivariate analyses to visualize and classify genome-wide gene expression data, recently reviewed in [28], have become the mainstream tool of single-cell RNA-seq, where they are used to discover cell types and quantify differences between them. With appropriate experimental design, in OMOs these analyses can lead to much more definitive biological conclusions than studies scrutinizing long lists of differentially expressed genes passing a certain significance cutoff.

**Table 2.**
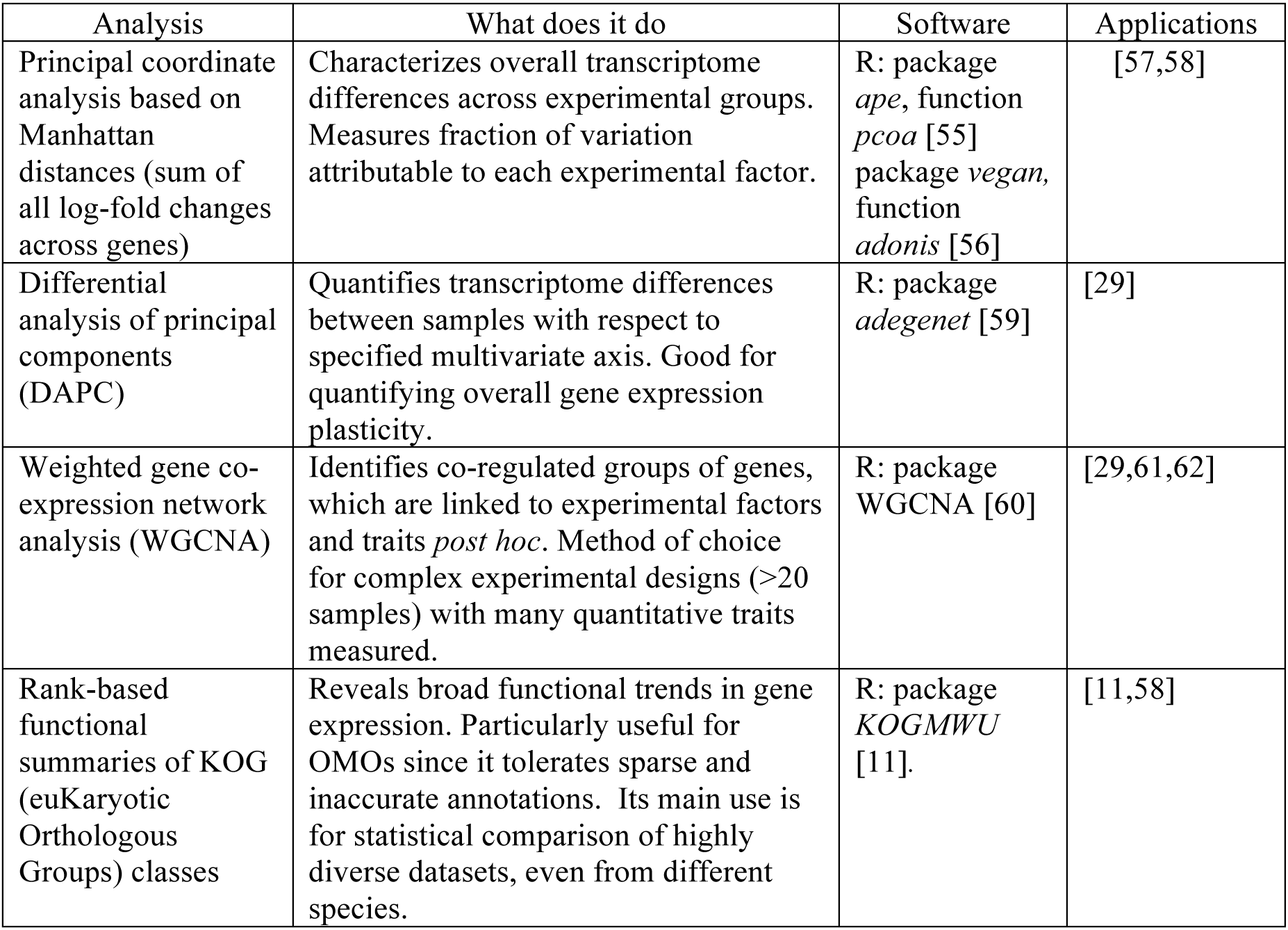
**Gene expression analyses not relying on accurate gene annotations**

### Gene expression as functional summary of genotype

Gene expression is best known for its context-dependence reflecting phenotypic plasticity, which is the view inherited from biomedical research dealing with genetically uniform models. In natural populations, one of the most important sources of gene expression variation is genetic difference among individuals, manifested as context-*in*dependent, individual-specific deviations in gene expression. This is easy to demonstrate in clonally replicated organisms such as corals. In two reciprocal transplantation experiments performed on different coral species in different oceans, stable between-genotype differences accounted for more than 50% of total gene expression variation despite transplantation of clonal fragments for up to a year to highly dissimilar sites [29,30]. In non-clonal model organisms such as mice or humans, the best demonstration of the effect of genetic variation on gene expression are abundant differences in expression between alleles of the same gene [31,32]. In humans, fixed between-population differences are exemplified by hundreds of genes that are differentially expressed between African and European Americans [33]. All this suggests that gene expression can be a proxy of not only phenotypic plasticity and acclimatization, but of genetic variation and adaptation. A major advantage of the use of gene expression for these types of studies is that gene expression integrates over many functionally relevant variants in the genome and thus represents a condensed functional summary of the genotype [34].

In humans, nearly half of all genetic variants affecting gene expression have detectable effects in all tissues [32], and so one feasible way to separate genotype-specific gene expression from context-dependent variation might be to perform “cross-tissue” comparison (see Glossary) to isolate body-wide expression shifts [35]. In the coming years, cross-tissue or similar analysis is likely to become a major approach to study functional genetic variation in natural populations.

## Epigenetics

Among many covalent chromatin modifications I will discuss DNA methylation since it currently receives the most attention in OMOs. Still it must be mentioned that in plants histone methylation appears be no less and perhaps even more involved in acclimatization and transgenerational plasticity [36]. While vertebrates show high methylation throughout the genome, invertebrates and plants methylate their genomes sparsely and mostly in protein-coding regions (so-called gene body methylation, GBM, [37]). The function of his ubiquitous and evolutionarily ancient DNA modification remains unclear [38,39] and the greatest challenge in the next few years will be to decipher it. The most important questions are: (i) Does GBM affect gene expression? (ii) Can it be modified on ecological timescale, to achieve acclimatization to a novel environment? (iii) Can acquired changes in GBM be transmitted across generations? If the answers to all three questions are “yes”, then we have a mechanism for transgenerational inheritance of acquired traits, which is an exciting (albeit tentative, [40]) possibility. Table 3 summarizes the methods for generating DNA methylation data. If every gene in the genome has to be interrogated, MBD-seq and meDIP provide the best resolution for sequencing effort [38]. If the goal of the study is to characterize general methylation patterns rather than identify specific genes, highly cost-efficient solutions are provided by RRBS-seq and methylRAD. For studies requiring single base resolution, the best approach appears to be direct detection by PacBio or ONT – however, these exciting developments still require validation in complex genomes.

**Table 3.** **Methods for interrogating DNA methylation**

| Method | Features | Pros | Cons |
| --- | --- | --- | --- |
| Whole-Genome Bisulfite Sequencing (WGBS) [63] | Sequences complete genome after bisulfite conversion | Complete characterization of 5me-cytosine methylation at single-base resolution | High coverage is required to obtain quantitative data. In non-vertebrate OMOs, much sequencing effort is wasted since most of genome is not methylated. |
| RRBS-seq [64] | Bisulfite sequencing of genome fragments adjacent to all (methylated and unmethylated) CCGG sites | Saves costs dramatically compared to WGBS. | Only a fraction of all CpG sites is interrogated. Complicated library preparation protocol. Sequencing effort is wasted on non-methylated sites. |
| MBD-seq [65], meDIP [66] | Pull-down and sequencing of methylated DNA. | Optimizes sequencing effort by focusing on methylated DNA. | Complicated library preparation protocol. Resolution equals the length of pulled-down fragments (~300-500b). Pull-down procedure is not absolutely efficient, many reads still correspond to un-methylated genome regions. |
| methylRAD [67] | Direct sequencing of genomic fragments adjacent only to the methylated CCGG and CCWGG sites. | Very simple library prep protocol. Highly cost-efficient due to focus on methylated sites only. | Only a fraction of all CpG sites is interrogated. New method, requires further benchmarking. |
| PacBio [68,69] | Direct detection of modified DNA bases during normal SMRT sequencing, based on polymerase lags. | Robust detection of 4-methylcytosine, 8-oxoguanine, and N6-methyladenine. Single-base resolution. | Same as WGBS. 5-methylcytosine, the most common methylation mark in animals, is not reliably detected. |
| ONT [70] | Direct detection of modified DNA bases during normal nanopore sequencing, based on conductivity changes. | Detects all marks, including 5-methylcytosine. Single-base resolution. | Same as WGBS. |

## Generating a reference sequence

For all approaches described here, the accuracy of the reference sequence in terms of per-base error rate must only be high enough to allow unambiguous mapping of high-accuracy (Illumina) reads. The gold standard of genome sequence quality, Q30 or 99.9% accuracy, would not provide any benefit compared to a rough draft accuracy of 99%. Occasional errors in the reference would manifest themselves as SNPs that are not polymorphic in the analyzed samples and therefore irrelevant for analysis. This is the same reason why it is possible to use a genome of a related species as a reference.

For AFS analysis, which does not require highly contiguous reference, even a rough genome draft that can be assembled from a single lane worth of 150b paired-end reads from Illumina HiSeq would be suitable. However, substantially better options are now becoming available for a comparably low price tag. The technology offered by 10x Genomics [41] attaches specific barcodes to short reads originating from the same long DNA fragment, which allows assembling Illumina HiSeq data into very long haplotypes. The two single-molecule long-read “third-generation sequencing” methods, Single Molecule Real Time (SMRT) sequencing by PacBio and nanopore sequencing by ONT, produce reads with broad length distribution, including exceedingly long ones (tens to hundreds of kilobases) resulting in a qualitatively more contiguous genome assemblies [42–45] (Table 4, see [43] for recent benchmarking study of assembly pipelines). At the moment of this writing, read accuracy and cost of data for PacBio (Sequel system) and ONT (R9 flow cell) were equivalent; PacBio generated higher proportion of long reads than ONT; however, PacBio’s library prep required ten fold more high-quality DNA than ONT. Both for PacBio and ONT it is critically important to obtain high molecular weight DNA in fragments exceeding 20kb in length. For new OMOs, it is also essential to confirm that the DNA is accessible to enzymatic modifications by trying to digest it with a frequent-cutting restriction endonuclease.

**Table 4.** **Assembly pipelines for PacBio and ONT reads**

| Pipeline | Required coverage | Features | Pros | Cons |
| --- | --- | --- | --- | --- |
| Canu + Quiver* [71] | >30x | Correct and trims reads before assembly. | Best accuracy at base, indel and assembly level. | Very computationally demanding for large genomes. Generates incomplete assemblies at low coverage. |
| Falcon + Quiver* [72] | >50x | Similar to Canu. | Standard for PacBio. | Very computationally demanding for large genomes. High reliance on reads >20kb. Highly incomplete assemblies at low coverage. |
| minimap + miniasm + racon [73,74] | <30x | Raw reads are assembled, correction is done post-assembly | Very fast even for large genomes. Works with lower coverage, shorter reads than Canu and Falcon. | The resulting accuracy is lower than with Canu + Quiver. |
| pilon [75] | NA (error correction method) | Performs additional correction post-assembly. | Boosts accuracy for any assembly. | Requires high-quality Illumina reads. |

\*Quiver is a consensus polishing software that is now replaced by Arrow to handle PacBio Sequel data (https://github.com/PacificBiosciences/GenomicConsensus). Racon [74] can be used instead of Quiver/Arrow [43].

For genome scanning, gene expression, or invertebrate DNA methylation analyses targeting protein-coding sequences (exome) genome sequence might not be the best reference; instead, a highly contiguous transcriptome assembly would be preferable. Until now the standard way to generate a *de novo* transcriptome was to perform high-coverage RNA-seq and assemble the results with Trinity [46]. In the coming years, it is expected that even higher-quality and lower-cost OMO transcriptomes would be generated by PacBio or ONT sequencing of full-length cDNA (or, for ONT, direct mRNA sequencing). The long-read capacity of these technologies would essentially obviate the need for assembly, leaving only the sequence correction procedure to be performed.

Finally, which tissue or body part to sample for sequencing? For genome sequencing it does not matter much as long as contamination by other DNA sources can be kept to a minimum, but for *de novo* transcriptomics it is not a trivial question, as gene expression varies dramatically across tissues and life cycle stages. In mammals, there is definitely an organ of choice that expresses nearly all genes in the genome: testis. Rather unexpected transcriptome complexity in the testis is putatively due to chromatin re-packaging during spermatogenesis, which results in genome-wide transcription leakage [47]. If so, testis might be a good choice for *de novo* transcriptomics not only for mammals but for any organism that produces compact sperm.

## Note on data sharing

As we have seen, the best power of ecological genomics in OMOs is achieved using a genome or transcriptome reference. Every new reference dataset enables new biological questions, and the whole OMO field will get a great boost if these resources are promptly shared. Please consider rapidly sharing your reference data, at least as soon as the initial preprint of your paper is posted to bioRxiv and ideally sooner, by distributing the link to data through research-related email list or professional twitter feed.

### Box 1: AFS models.

In the world of OMOs we are usually dealing with samples from many populations, which would be hard or impossible to model simultaneously; moreover, there are usually many populations left unsampled. To infer meaningful demographic parameters in a sparsely sampled system of many populations, a practical solution is to perform two-dimensional AFS analysis of all population pairs [10]. Typical hypotheses and corresponding tests are:

- Are the two populations demographically separate?

∘ compare model with split to model without split, under which the two compared populations are regarded as independent samples from the same population.
- If yes, is there still gene flow between them?

∘ compare split models with and without migration.
- If yes, is the gene flow symmetric or asymmetric?

∘ compare split model with two potentially different migration rates to a split model with a single symmetrical migration rate.
- Was population size stable or went through changes in the past?

∘ compare single-population model involving population size change in the past to a standard neutral model.

Simple command-line scripts for AFS plotting and running basic pairwise models in *moments* can be found here: https://github.com/z0on/AFS-analysis-with-moments. To access the full potential of *moments*, however, the user is expected to compose python scripts of their own.

